# Genome recombination-mediated tRNA up-regulation conducts general antibiotic resistance of bacteria at early stage

**DOI:** 10.1101/2021.01.19.427372

**Authors:** Huiying Fang, Guandi Zeng, Jing Zhao, Tingkai Zheng, Lina Xu, Wei Gu, Yutong Liu, Jinning Zhang, Xuesong Sun, Gong Zhang

**Affiliations:** Key Laboratory of Functional Protein Research of Guangdong Higher Education Institutes, Institute of Life and Health Engineering, Jinan University, Guangzhou 510632, China; State Key Laboratory of Biocontrol, College of Ecology and Evolution, Sun Yat-Sen University, Guangzhou 510275,China

## Abstract

Bacterial antibiotic resistance sets a great challenge to human health. It seems that the bacteria can spontaneously evolve resistance against any antibiotic within short time without the horizontal transfer of heterologous genes and before accumulating drug-resistant mutations. We have shown that the tRNA-mediated translational regulation counteracts the reactive oxygen species in bacteria. In this study, we demonstrated that isolated and subcultured *Escherichia coli* elevated its tRNAs under antibiotic stress to rapidly provide antibiotic resistance, especially at the early stage, before upregulating the efflux pump and evolving resistance mutations. The DNA recombination system repaired the antibiotic-induced DNA breakage in the genome, causing numerous structural variations. These structural variations are overrepresented near the tRNA genes, which indicated the cause of tRNA up-regulation. The strains knocking out the recombination system could not up-regulate tRNAs, and coincidently, they could hardly evolve antibiotic resistance in multiple antibiotics, respectively. With these results, we proposed a multi-stage model of bacterial antibiotic resistance in an isolated scenario: the early stage (recombination – tRNA up-regulation – translational regulation); the medium stage (up-regulation of efflux pump); the late stage (resistance mutations). These results also indicated that the bacterial DNA recombination system and tRNA could be targeted to retard the bacterial spontaneous drug resistance.

## Introduction

Bacterial antibiotic resistance (AR) is a major threat on health. The number of newly developed antibiotics is rapidly declining over years [1]. In sharp contrast, bacteria evolve AR shortly after the application of newly developed antibiotics [2], emphasizing an alarming “antibiotic crisis” [3, 4]. Studies have shown that numerous AR genes exist in nature for centuries already, providing AR by horizontal transfer [5–7]. However, this theory is difficult to explain the fact that bacteria quickly evolve resistance against non-natural, fully artificially synthesized antibiotics [2], for which the specific natural AR genes rarely exist. This indicates a general and intrinsic mechanism of bacteria to resist any antibiotics without any heterologous AR gene input.

Two major types of spontaneous AR mechanisms has been revealed without any heterologous AR gene input: (a) mutation or modification of the antibiotic targets; (b) increasing efflux or reducing membrane permeability to limit the cellular antibiotic concentration [8]. However, these mechanisms take effect quite slowly. For example, continuous antibiotic pressure leads to accumulation of genomic point mutations of *Escherichia coli* starting from 2~5 days, causing non-synonymous mutations and thus alters the proteins, spontaneously evolving AR without heterogenous genes [9]. If the bacteria cannot withstand the damage caused by the drugs at the beginning, they have no chance to develop specific resistance (e.g. genetic mutations). These facts indicate that bacteria have an intrinsic and general mechanism to evolve early AR independent of point mutations or specific AR genes.

It is known that most antibiotics facilitate generation of reactive oxygen species (ROS) in cells, which damages nucleic acids and proteins. Therefore, counteracting ROS is a must to resist most antibiotics. Therefore, we posit that the abovementioned the tRNA regulation as intrinsic and general mechanism, for early AR. Revealing such mechanism would help us to understand the robustness of bacteria and find an alternative solution against AR.

## Materials and Methods

### Bacterial strains

*Escherichia coli* (*E.coli*)strains BW25113 and BW26355 (ΔrecA version of BW25113) was purchased from the Coli Genetic Stock Center of the Yale University. The reference genome sequence NZ_CP009273.1 and its annotation (downloaded from NCBI) were used in the bioinformatics.

### Subculturing bacteria in antibiotics

The sensitive bacteria (the BW25113 strain) was cultured in the LB medium from a single colony as the generation 0. A new generation of culture was inoculated at 1:500 ratio and cultured at 37°C for 12 hours with the presence of the antibiotics at 1/2 minimal inhibition concentration (MIC). The MIC was measured for each generation.

### Competition assay

The *E. coli* BW25113 transformed with pBAD33 and pRIL plasmid, respectively, were mixed at 1:1 ratio. The mixture was subcultured in LB medium with the presence of 100 μg/mL chloramphenicol and ciprofloxacin at 1/2 MIC. As a control, the mixture was subcultured in LB medium with the presence of chloramphenicol but without ciprofloxacin. For each generation, the plasmids were extracted from the bacterial culture and resolved in agarose gels.

### Whole genome sequencing

The genomic DNA was extracted using HiPure Bacterial DNA extraction kit (Magen). The genomic DNA was sonicated into ~300bp fragments. The whole-genome sequencing library was constructed using MGIEasy DNA library construction kit (MGI) following the manufacturer’s instructions. The libraries were sequenced on an MGISEQ-2000 sequencer at PE100 mode. The raw sequencing data was deposited to SRA database (accession: PRJNA644774).

For single nucleotide variation (SNV) analysis, the reads were mapped to reference genome using FANSe3 with the parameters -E3 -S13 --indel. The SNV were detected using the previously published and experimentally validated method [10].

For structural variation (SV) analysis, each read was trimmed into 26nt reads. To balance the throughput of each generation, 57.4M reads were used in the dataset of each generation. The reads were mapped to the reference genome using FANSe3 with the parameters -E3 -S13 --unique. The entire genome was divided into 600 nt bins. The uniquely mapped reads were merged according to the bins. The reads, whose two ends were mapped far away (more than 1000 nt apart) or mapped to the same direction (indicating inversion) were counted as “SV reads”. Fisher exact test was performed for each bin with the null hypothesis that no SV happens.

Nanopore single-molecule long-read sequencing was performed according to the manufacturer’s protocol. Base calling was conducted using Guppy software with default parameters. The raw reads data was deposited to SRA database (accession: SRR12717711). Reads were aligned to BW25113 reference genome using Minimap2 software. Structural Variations were called using SVIM software with default parameters.

### Transcriptome sequencing

The bacterial cells were harvested by centrifugation at 4°C 4000 xg for 5 minutes. The cell pellet was resuspended and washed using PBS. The cells were treated using 1.25 mg/mL lysozyme at 4°C for 10 min and collected by centrifugation at 5000 xg for 5 min. The pellet was dissolved in 1 mL Trizol reagent and the RNA was extracted using Trizol method. The rRNA was removed using the RiboX rRNA removal kit (Chi-Biotech) following the instructions. The RNA library was constructed using the MGIEasy mRNA library prep kit V2 and sequenced on an BGISEQ-500 sequencer at SE50 mode. The raw sequencing data was deposited to SRA database (accession: PRJNA644774).

The gene expression levels were quantified using the Chi-Cloud NGS analysis platform (http://www.chi-biotech.com/technology.html?ty=ypt). In brief, the reads were mapped to the reference genome using FANSe3 with the parameters -E3 --indel. The gene expression was measured using rpkM method. The log10 rpkM values were subjected to the correlation, PCA and clustering analyses. Gene Ontology overrepresentation analysis were performed using PANTHER-DB (http://www.pantherdb.org/). Significance was considered when FDR<0.01.

### tRNA sequencing

We developed a method to specifically sequence tRNAs in BGISEQ/MGISEQ sequencers. The sequences of adapters and primers are: Y5: 5′-TAAGACCGCTTGGCCTCCGACTTACTGGATACTGG+rN (rN = equal mixture of rA, rT, rG, rC), Y3: 5′-GTATCCAGTN_16_AAGTCGGATCGTAGCCATG (N_16_ = 16 random nucleotides), barcode primer: 5′-TGTGAGCCAAGGAGTTGTAGT GGGGATTTGTCTTCCTAAGACCGCTTGGCCTCCGACT, adapter primer: 5′-GAACGACATGGCTACGATCCGACTT (5′-phosphorylated). These adapters and primers were mixed at 1:1 ratio. Starting from 6μg total RNA of *E. coli*, 60ng human tRNA was added as spike-in. The tRNAs were deaminoacylated in 0.2M pH=9.5 Tris-HCl buffer at 37°C for 45 min. The RNA was then purified using HiPure RNA pure micro kits (Magen). 200~300ng RNA was mixed with the primers. Ligation was conducted using T4 double stranded RNA ligase 2 (NEB) following the manufacturer’s instructions. Reverse transcription was performed using the SuperScript III Reverse Transcriptase (Invitrogen) at 55°C for 45 min and 70°C for 15 min. The library was PCR amplified using the barcode primer and the adapter primer for 20 cycles. The library was sequenced on a BGISEQ-500 sequencer at SE100 mode. The raw sequencing data was deposited to SRA database (accession: PRJNA644774).

The *E. coli* and human tRNA reference sequences were downloaded from the gtRNAdb (http://gtrnadb.ucsc.edu/). The reads were mapped to the reference sequences using BLAST (run in local server). The tRNA read counts were normalized according to the spike-in tRNA read counts.

## Results

### Bacteria evolve antibiotic resistance without detectable mutations in protein-coding regions

We cultured *E. coli* BW25113 in LB minimal medium under ciprofloxacin (CIP) at a concentration of 1/2 MIC (minimal inhibition concentration) in a continuous subculture way. New generation of culture is inoculated subsequently from previous culture every 12 hours. In the entire process, *E. coli* was cultured in sterile flasks to avoid heterogeneous genes. The MIC gradually increased over 330-folds after 45 generation subculture (Fig. 1A). The growth rates decreased during the subculturing and almost reached constant after the 24^th^ generation (Fig. 1B). Visually, these phenotypes divide the entire subculturing into three distinct stages: (a) the early stage (before the 7^th^ generation), where the MIC instantly increased but fluctuate around 0.1~0.2 μg/ml, and the growth rate remarkably decreased; (b) the medium stage (approximately generation 7~21), where the MIC steadily increased, and the growth rate steadily decreased; (c) the late stage (from the 22^nd^ generation), where the MIC increased in a zigzagged manner to more than 100-folds compared to the sensitive strain, and the growth rate maintained at an almost constant level, less than half of which of the sensitive strain. These three stages are marked on the Fig.1A~B. The stage-wise evolution of the AR indicated that the bacteria respond to the CIP using distinct mechanisms in each stage.

**Fig. 1:**
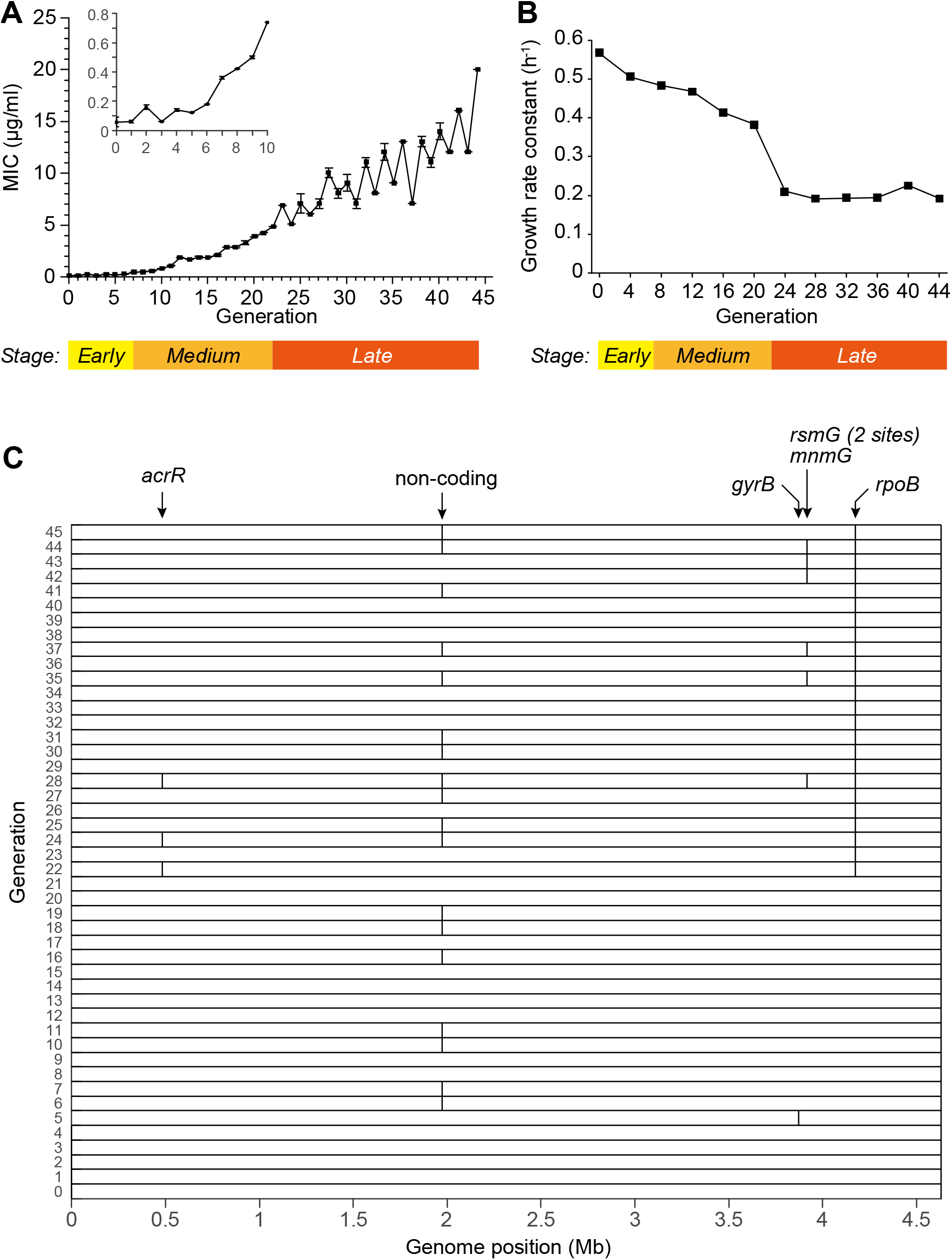
Subculturing *E. coli* BW25113 in the sub-lethal concentration of CIP. (A) The MIC of each generation. (B) The growth rate constant measured every 4 generations. (C) Single nucleotide mutations detected in the WGS of each generation, marked as vertical bars. The affected genes were marked on the top. Some closely-located mutations are not distinguishable in the figure due to the limited resolution. Detailed information of each mutation is listed in the Supplementary Table S2.

We first speculated that the AR was endowed by mutations in genes. To screen the non-synonymous mutations that may associate with the spontaneous drug resistance, we performed whole genome sequencing (WGS) for each generation, yielding more than 200x sequencing depths and more than 98.98% coverage (Supplementary Table S1). SNV calling were performed according to the published method, whose accuracy and sensitivity were validated via massive Sanger sequencing [10]. Only 7 SNVs were identified from the subcultured bacteria, and 6 of them were in the CDS (Fig. 1C, Supplementary Table S2). Among them, only one mutation in the gene *rpoB* were persistently observed from the 22^nd^ generation, which is known to endow resistance [11]. However, all the other mutations were sporadically detected in a few generations. For example, the drug target of CIP, the *gyrB* gene, was found mutated only in the 5th generation and not in any other generations. Thus, this mutation is unlikely to explain the resistance. This result showed that no meaningful mutations in coding genes were detected in the early stages.

### Translation is up-regulated as a response against antibiotics at early stage

Since no mutations were associated with the AR at early stages, we next performed RNA-seq for each generation to analyze the temporal change of the transcriptome in the presence of CIP. The PCA plot of the gene expression levels can be clustered into three distinct stages: the early stage (original and generations 1~7), the medium stage (generations 8~21) and the late stage (generations 22~45) (Fig. 2A). The mutual Pearson correlation matrix also confirmed this (Fig. 2B). This correspond to the three stages illustrated in Fig. 1A~B.

**Fig. 2:**
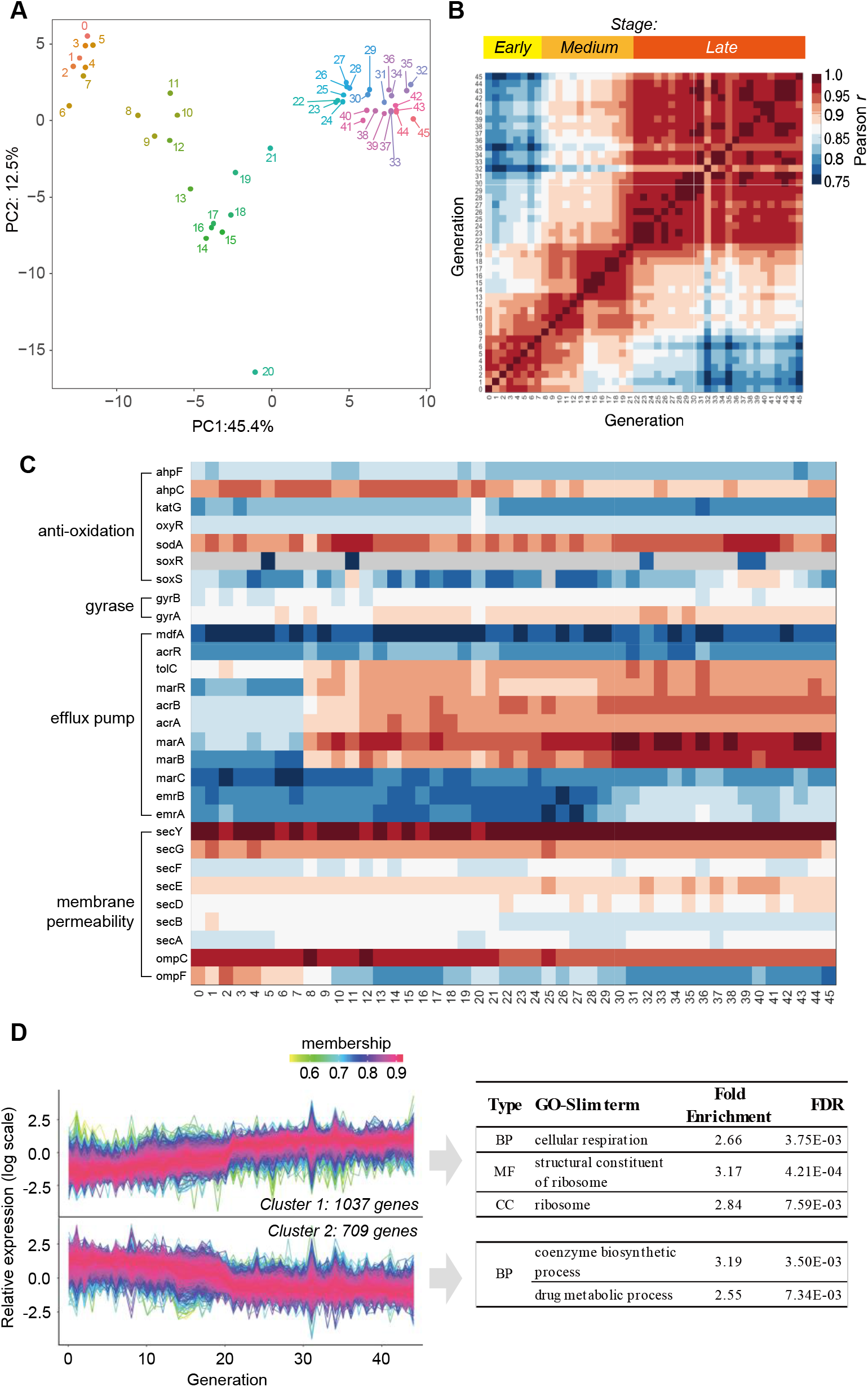
Transcriptome changes during the subculture. (A) PCA analysis of the gene expression profiles of each generation. Numbers denote the generation. (B) Mutual Pearson correlation coefficients of log RPKM values between each two generations. (C) Temporal change of gene expression of genes that are involved in traditional AR mechanisms. (D) Expressed genes were clustered into two clusters according to the expression trend over the generations. The GO-Slim terms with fold enrichment > 2 and FDR < 0.01 were filtered and listed.

We then looked deeper into the temporal expression profiles of traditional resistance genes (Fig. 2C). Under the constant presence of CIP, which induces cellular ROS, the antioxidation system SoxR and SoxS did not increase over the generations. The DNA gyrases, which are the target of CIP, also did not change significantly over the generations. The only obvious trend is the elevated expression of *acrA*, *acrB*, *marA*, *marB* and *marR*, which are associated with drug efflux. Nevertheless, these genes did not overexpress before the 8^th^ generation, indicating that they are not the cause of AR at the early stage. Moreover, the genes regulating outer membrane permeability and most drug efflux genes (e.g. *marC*, *emrA*, *emrB*, *lamb*, *phoE*, etc.) were not up-regulated implying drug uptake and efflux was not altered. These results showed that the traditional resistance mechanisms cannot explain the early-stage response.

Among the 2429 genes which were quantified in all generations, 1746 genes showed significant trend over the generations (*P*<0.05, Mann-Kendall trend test). These genes can be clustered into two categories: the gradually up-regulated and down-regulated genes (Fig. 2D). Increasing the number of clusters to 4 revealed the same trend (Supplementary Fig. S1), showing the robustness of the clustering analysis. The GO-Slim enrichment analysis showed that the gradually up-regulated genes were mainly enriched in respiration (energy production) and ribosome, while the coenzyme biosynthetic process and drug metabolism process. This result hinted that the translation may be enhanced as the major response to the antibiotics.

### tRNA up-regulation during the subculture provided antibiotic resistance

Since the ribosomes were constantly up-regulated during the subculture, it is logic to assume that the tRNAs should be also up-regulated to maintain efficient translation. To test this postulation, we loaded equal amount of total RNA (not from the equal number of cells) of each generation on gels. On agarose gel, the 23S and 16S rRNA bands were equal in all generations. However, the 5S rRNA + tRNA band gradually increased till the 35^th^ generation (Fig. 3A). On polyacrylamide gel, which resolved 5S rRNA and tRNAs, the 5S rRNA were equal in all generations, and the tRNA bands increased (Fig. 3B). To investigate the up-regulation of tRNAs, we then performed tRNA-seq using artificially synthesized tRNA as spike-in to quantify the tRNA levels. The tRNA reads were normalized against the spike-in. The result showed that the total tRNA increased during the entire subculture, even at the early stage (Fig. 3C), which reproduced the trend on gels. Moreover, most tRNA species were up-regulated at early stage (Fig. 3D), indicating that many tRNAs were affected. In contrast, the translation elongation factors (*tufA/tufB* for EF-Tu and *tsf* for EF-Ts) and the ribosomal proteins (e.g. *rplA* for large ribosomal unit and *rpsA* for small ribosomal unit) were not increased over the entire subculturing, at least on transcription level (Fig. 3E). Together with the constant rRNA content (Fig. 3A~B), these results demonstrated that the translational machinery was not up-regulated. Therefore, tRNA up-regulation would effectively maintain the translational elongation and thus help the survival under the ROS induced by CIP at early stage.

**Fig. 3:**
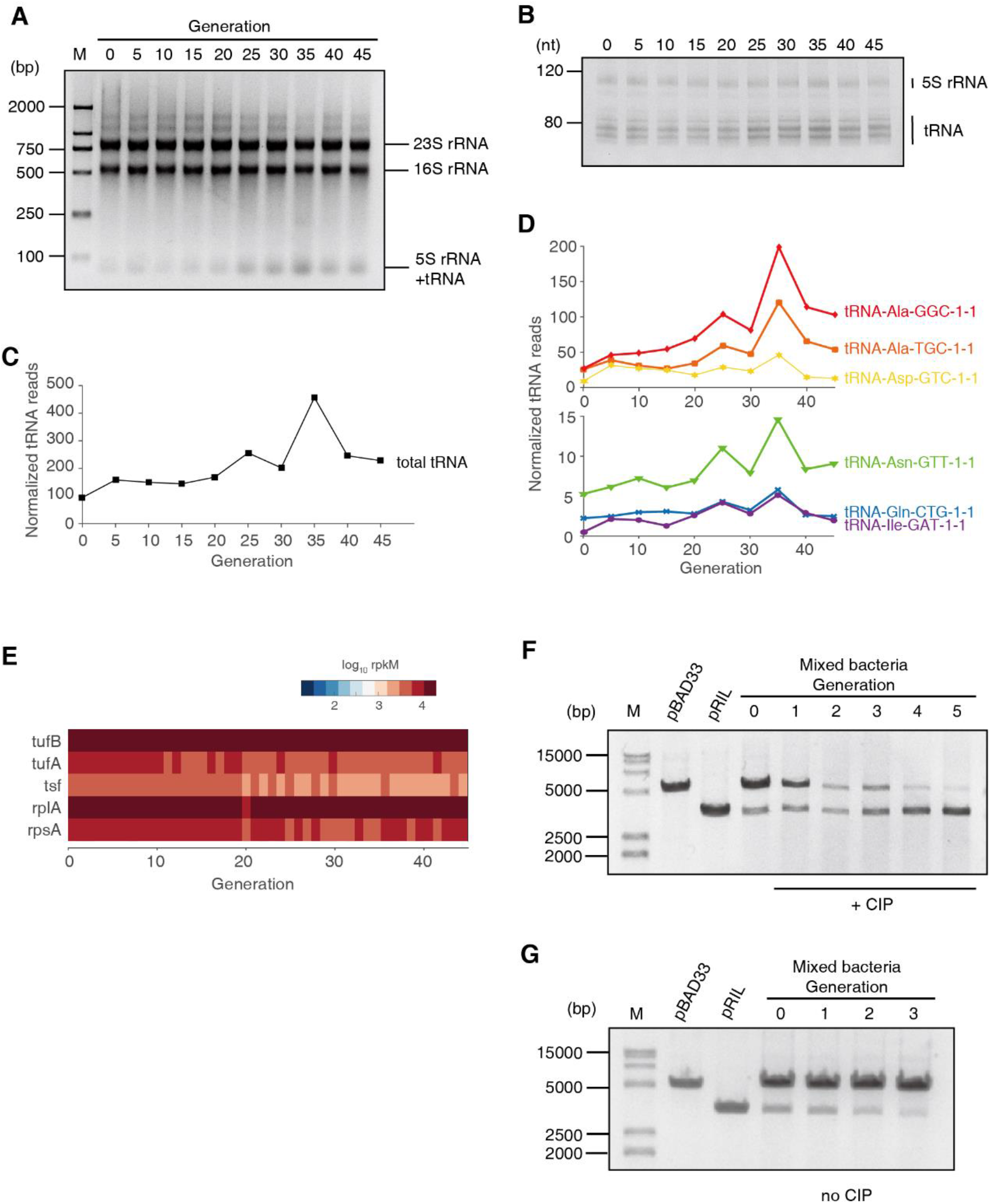
tRNA up-regulation under the CIP stress. (A) Agarose gel of total RNA of each generation. Equal amount of total RNA was loaded. (B) Polyacrylamide gel of total RNA of each generation. (C) Total tRNA abundance of each generation, quantified using tRNA-seq, normalized using artificially synthesized tRNA spike-in. (D) The abundance of 6 most abundant tRNAs during the subculturing, normalized using the spike-in. (E) The expression of elongation factors and ribosomal proteins. (F,G) The competition assay. pBAD33 and pRIL plasmids were transformed into *E. coli* BW25113, respectively, and mixed to start subculturing. For each generation, the plasmids were extracted from the mixture. (F) Subculture with CIP. (G) Subculture without CIP.

To further examine the significance of tRNA upregulation at early stages of AR, we performed the competition experiment. We transformed the pRIL (which expresses three tRNAs) and pBAD33 (which does not contain any tRNA, serving as a negative control) plasmids into *E. coli* BW25113 strain, respectively. This pair of plasmids were used in previous studies to investigate the effect of tRNA up-regulation [12]. We mixed these two types of bacteria together in the same tube and started subculturing with and without sub-lethal concentration of CIP respectively. For each generation (12 h culture), the plasmids of the mixed bacteria were extracted and resolved on agarose gel. With CIP culture, the pRIL plasmid became dominating after 3 generations, and the pBAD33 almost diminished after 5 generations, demonstrating that the excessive tRNA provides growth advantage under antibiotics (Fig. 3E). In contrast, without CIP, the pRIL-containing bacteria cannot compete with the pBAD33-containing counterparts, and the pRIL plasmid almost diminished after 3 generations (Fig. 3F). This coincides with our previous studies: under the normal conditions, the excessive tRNAs will suppress translational pausing and thus lead to massive protein misfolding and aggregation [12–14]. Under the ROS (generated by CIP), the translation elongation is globally decelerated, and thus the translational pausing is not an issue. Up-regulated tRNAs will maintain the protein production to counteract the oxidative lesion [12]. In sum, these results demonstrated that tRNA up-regulation provided survival competence under the antibiotics by maintaining translation activity.

### Genome recombination is enriched near tRNA genes

Next, we need to explain the potential reason of tRNA up-regulation upon CIP pressure. We first checked the expression levels of the transcription factors of tRNA genes. However, the transcription factor *fis* for tRNA genes and the other major transcription factors were not increased (Fig. 4B). The *E. coli* tRNA genes are transcribed in operons, started from promotors. However, no point mutation was discovered in or near the tRNA genes. Nevertheless, point mutation is not necessary for tRNA up-regulation. If fused into other operons, the tRNA gene will be expressed under the control of that operon, and can be expressed more than 10-folds than the wild-type, which has been observed in literature [15]. CIP kills bacteria by causing DNA double-strand breakage via producing ROS and targeting GyrA/GyrB. Such DNA lesion will be repaired primarily by recombination system. Therefore, we postulate that genome recombination happened near tRNA genes to interfere tRNA expression.

**Fig. 4:**
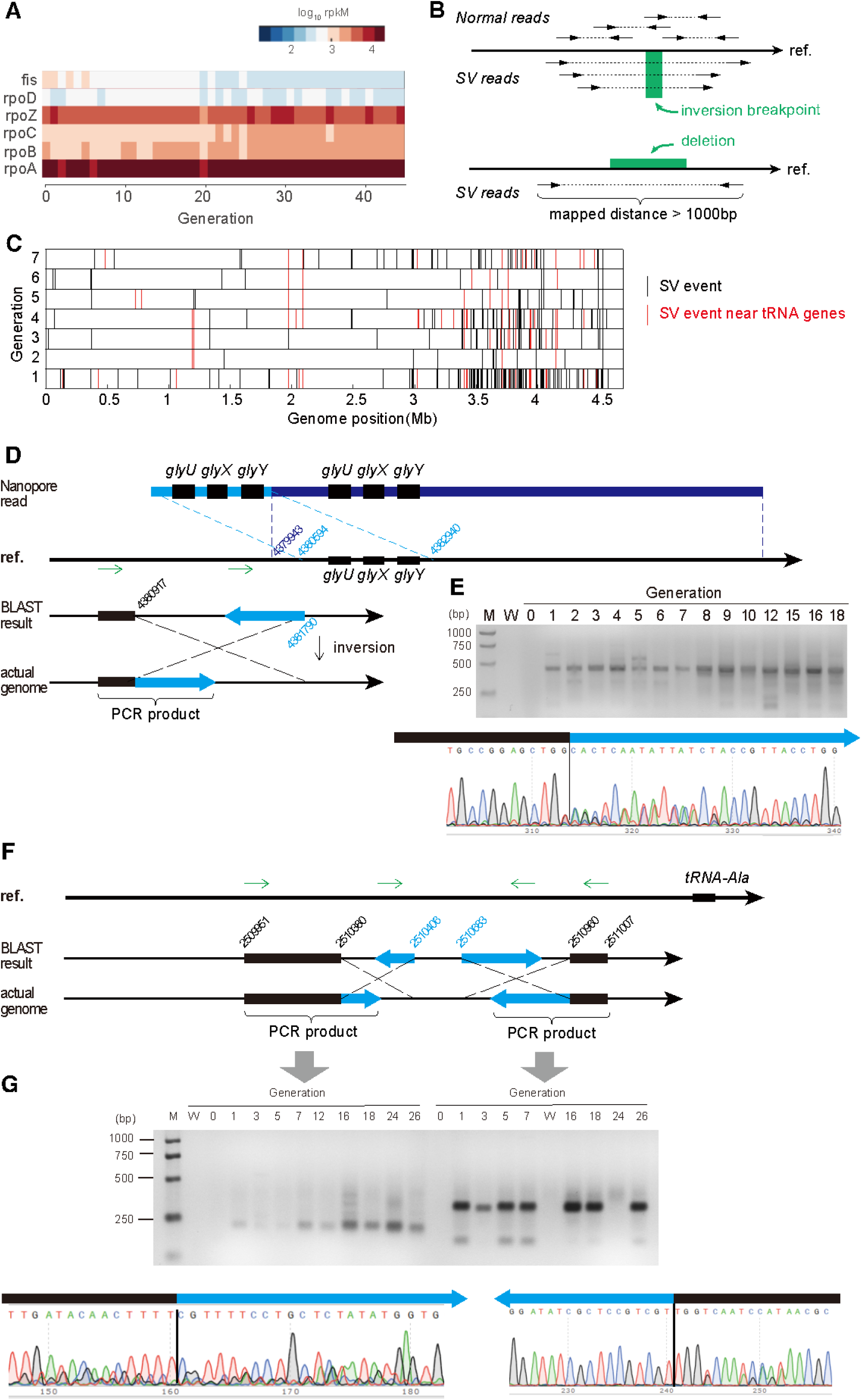
The SV that affects the tRNAs. (A) The expression levels of the major transcription factors that are involved in tRNA expression. (B) Detecting SV events using paired-end reads. The WGS library was constructed as ~300bp insert. After mapped to the reference genome sequence (“ref.”), the two ends of the same read should be mapped in opposite direction. An inversion breakpoint candidate is identified if the two ends of the same read are mapped in the same direction. Deletion of a fragment is detected if the two ends of the same read are mapped in opposite direction but far apart. Only the uniquely mapped reads were taken into consideration. (C) The SV events detected using WGS data during the early stage. Red bars represent the SV events near the tRNA genes, and the black bars represent the other events. (D) An example of a duplication event and an inversion event near *glyU*-*glyX*-*glyY* operon. (E) The gel electrophoresis of the PCR product validating the inversion event in panel D. The PCR was performed using the two primers that are indicated as green arrows in panel D. Lane M = marker; lane W = water control (negative control using water as template). The Sanger sequencing of the PCR product is exhibited for the junction of the inversion. (F) Two inversion events near the tRNA-Ala gene. (G) The gel electrophoresis of the PCR product validating the inversion events in panel F. The PCR was performed using the primers that are indicated as green arrows in panel F. Lane M = marker; lane W = water control (negative control using water as template). The Sanger sequencing of the PCR products are exhibited for the junctions of the inversion.

To test this hypothesis, we analyzed the structural variation (SV) of the bacterial genome from the WGS paired-end sequencing data (Fig. 4B). From the first generation after applying CIP, many SVs were detected, including deletions and inversions (Fig. 4C). In the first 7 generations, 341 SVs were detected, and 63 SVs were detected near the tRNA genes (defined as +/− 3000 nt near tRNA genes). The actual probability that an SV occurs near tRNA genes was significantly higher than the random distribution (*P* = 4.74×10^−35^, Fisher Exact Test), suggesting that the tRNA genes and their flanking sequences are more susceptible to the double-strand break induced by antibiotics. To avoid the limitations and bias of the short-read next-generation sequencing, we subjected the genomic DNA to Nanopore sequencing to obtain long reads. We found 1408 SV events in the nanopore sequencing dataset, and 57 among them were located near the tRNA genes (*P*=8.97×10^−6^, Fisher exact test). This also concluded that the SV events are significantly overrepresented near the tRNA genes, which coincides to the MGISEQ-2000 WGS conclusion. We found a long read that contained two sets of glycine tRNAs *(glyU-glyX-glyY*) (Fig. 4D). This 23394 nt read was aligned to reference genome as two segments, demonstrating a duplication that multiplicated tRNA copy number.

An inversion event was also found in the upstream of the *glyU-glyX-glyY* operon at 4380917 supported by 21 paired-end reads, whose both ends were mapped to the reference genome in the same direction (Supplementary Fig. 2A). Using two primers in the same direction (designed according to the reference genome), no PCR result could be yielded unless an inversion occurs (Fig. 4D). Indeed, a band of 415 bp PCR product was found (Fig. 4E). After Sanger sequencing, the fragment can be aligned to the reference genome in two segments in opposite direction, demonstrating an inversion from 4380917 to 4381790 (Fig. 4D~E). Coincidently, the tRNA-Gly-GCC, which is transcribed from the *glyU-glyX-glyY* operon, was found increased in the ciprofloxacin stress (Supplementary Fig. S3). Two inversions were detected in the upstream sequence of tRNA-Ala: 2510380-2510406 and 2510683-2510960 (Fig. 4F). These events were supported by 18 and 11 paired-end reads with the same mapped direction of both ends (Supplementary Fig. S2B). These two events were also validated using the abovementioned PCR method and Sanger sequencing (Fig. 4G). Coincidently, the alanine tRNA was upregulated in the ciprofloxacin stress (Fig. 3D). Notably, all these PCR evidence showed that these SV events were absent in the original sensitive strain (generation 0), while constantly appear since the first generation under ciprofloxacin stress, indicating that these SV events were induced by antibiotics.

The tRNA-processing enzymes may also be affected by the SV events. For example, a deletion happened between 3916925 to 3917502, which was detected from the 21^st^ generation, was confirmed by Sanger sequencing (Supplementary Fig. S4A) and PCR (Supplementary Fig. S4B). Accordingly, the expression of *mnmG* gene was remarkably elevated since the 21^st^ generation (Supplementary Fig. S4C). This deletion causes a truncation of the *mnmG* gene. However, it seems that this truncation does not affect the catalytic domain of MnmG enzyme. The nucleotide binding sites of MnmG are all before 370 aa, which is in its main domain, according to the UniProtKB (Supplementary Fig. S4D, blue domain). However, the truncation cuts the fragment 497-629aa, which is in a flexibly-linked independent domain 479-629aa (Supplementary Fig. S4D, orange domain). Therefore, the truncation remained an intact main domain, which should maintain its catalytic function. The MnmG is a 5-carboxymethylaminomethyluridine-tRNA synthase. It is responsible for tRNA modification that reduces frameshift errors in translation, according to the EcoCyc database. Therefore, the SV related to the up-regulation of MnmG may consolidate the protein synthesis quality, which facilitates the stress response.

### Suppressing recombination repair system decreases the evolution of intrinsic AR

To validate the hypothesis that the recombinant repair causes the tRNA upregulation and thus leads to AR, we suppressed the recombination enzymes by gene knockout. The *E. coli* BW26355 is the Δ*recA* strain (*recA* gene was knocked out from the *E. coli* BW25113 strain). We also cloned the *recA* gene into pET-28b plasmid and transferred to the BW26355 strain to restore RecA expression to the wild-type level (Fig. 5A). We subjected these three bacteria to subculture under sub-lethal level of CIP. As expected, the Δ*rec*A strain evolved AR much slower than the wild-type and the *recA* restored strain. The final MIC that the Δ*recA* strain could reach is also much lower than the other two strains with RecA (Fig. 5B). The tRNA content of the Δ*recA* strain did not show a visible increase (Fig. 5C). Similar trend also applied to other kinds of antibiotics, such as gentamycin (Fig. 5D) and ampicillin (Fig. 5E), under which stress Δ*recA* strain evolved AR slower, and showed the maximum MIC lower than the wild-type.

**Fig. 5:**
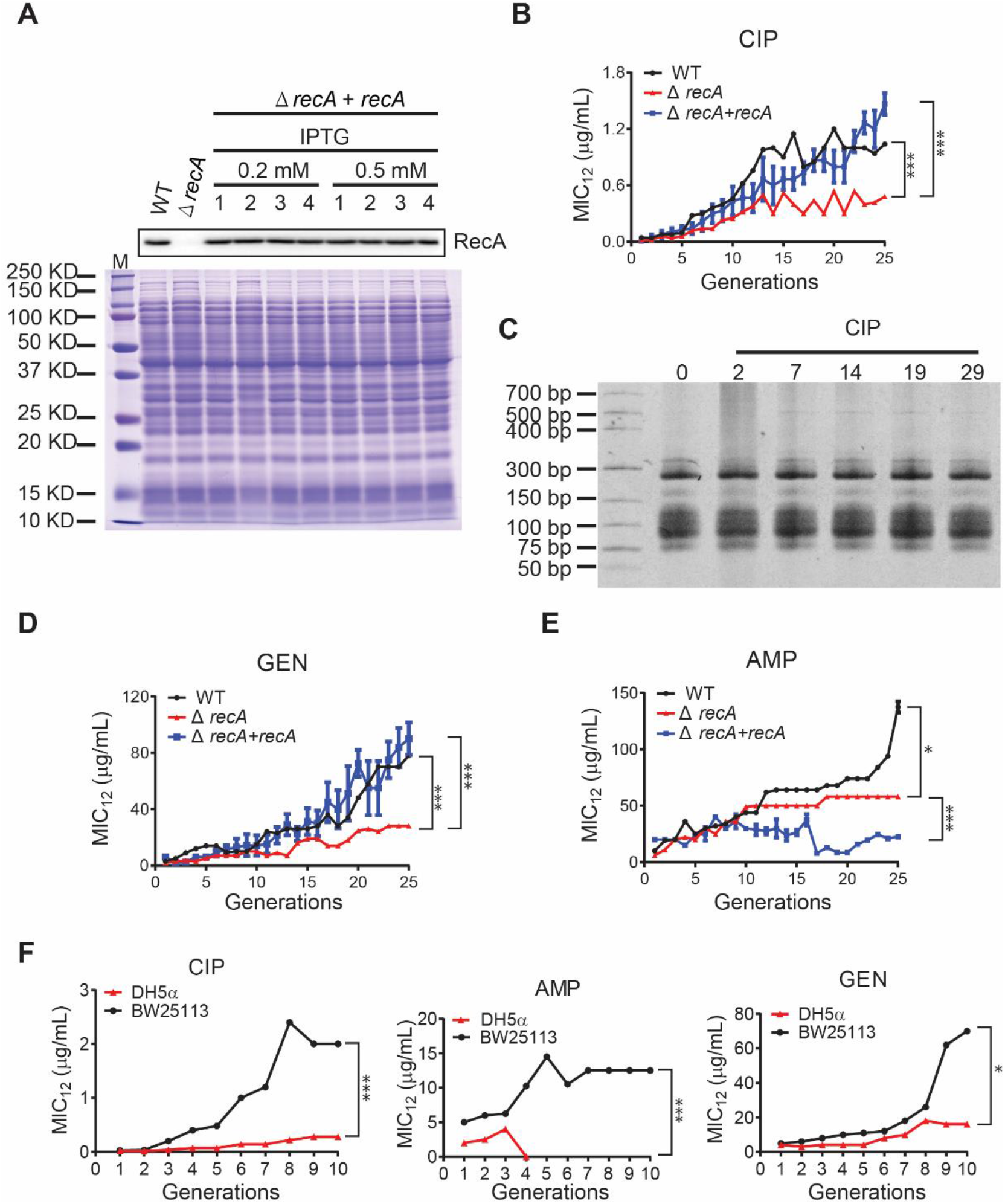
Repressing the recombination abolish the AR evolution. (A) The deletion of the *recA* gene (*ΔrecA* strain) and the RecA-restored strain (Δ*recA* + *recA*). Western blot confirmed the RecA expression levels. The total protein gel stained with Coomassie blue was used as loading control for the same loading. 1~4 represent the 4 independent biological replicates. (B) The MIC of the WT (black line), Δ*recA* (red line) and RecA-restored (blue line) strains, respectively, during the subculturing with CIP. (C) The tRNA content during the subculturing with CIP, resolved in a PAGE. (D,E) The MIC of the three strains during the subculturing with gentamycin (GEN, panel D) and ampicillin (AMP, panel E), respectively. (F) The MIC of the WT (black line) and the DH5α strain (red line) during the subculturing with CIP, AMP and GEN, respectively. All MICs were illustrated as mean±SD of three biological replicates. *, *P <* 0.05. ***, *P* < 0.0001.

The *E. coli* SOS repair system include four components, namely RecA, LexA, RelA and EndA. RecA is the most important one, but the other three are not negligible. The *E. coli* DH5α strain lacks all these four recombinases. Therefore, the DH5α strain has minimum recombination ability, much lower than the Δ*recA* strain. Indeed, when subcultured, DH5α can hardly evolve AR (Fig. 5F). In the ampicillin-containing media, the DH5α die out after 3 generations. These results showed that the recombination ability is positively correlated to the AR evolution, and this applies to multiple antibiotics.

## Discussion

Previous studies on intrinsic AR mainly focused on the mutations, membrane permeability and the efflux pump up-regulation. However, these three mechanisms need time to accumulate and may need days to evolve [9, 16, 17]. The early stage of the AR evolution, especially the general resistance mechanism against almost all kinds of antibiotics, has been overlooked. Such mechanisms are essential for bacteria to promptly combat the initial attack of antibiotics and thus make time for more specific and efficient resistance to evolve. Here, we provided an evidence that the translational response can serve as a general mechanism for AR at early stage. Since most antibiotics induce ROS in bacteria [18], which induces DNA damage, bacteria start the recombinant repair to response ROS stress which may lead to massive recombination in genomes. The recombination events are enriched near the tRNA genes in bacteria under the constant CIP pressure, probably due to the highly homologous sequences [19, 20]. This will, by chance, elevate the tRNA expression, and thus maintain the translational efficiency to synthesize enough proteins to maintain physiological processes, fix the impaired components and counteract the stress [12], which is essential for the survival [21], especially at the early stage, and thus protect the cell from the destruction (Fig. 6A).

**Fig. 6:**
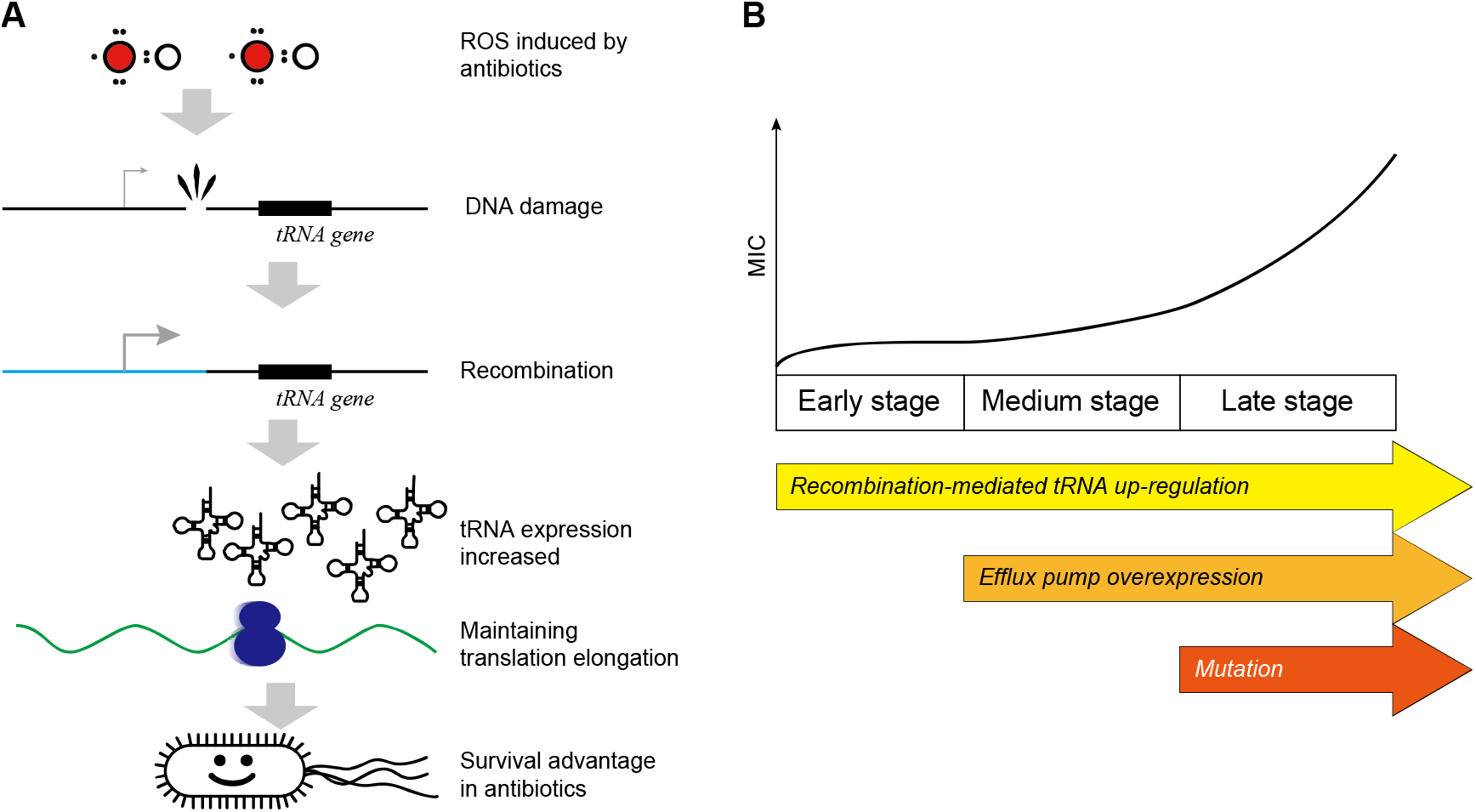
Scheme of the recombination-mediated tRNA up-regulation. (A) Pathway illustration of the recombination-mediated tRNA up-regulation. (B) Multiple AR mechanisms coordinate in a step-wise manner.

As a general mechanism, the recombination and tRNA-mediated general AR at early stage does not target specific antibiotics, but rather apply to the antibiotics which induce ROS. It can be explained that the ROS is a major cause of DNA damage and thus induces the recombination. The antibiotics CIP, GEN, AMP induce ROS in different extent, and thus can be counteracted by tRNA up-regulation. The antibiotics used in Hoeskema *et al.* (AMX, ENRO, KAN, TET) belongs also to the same category, and they observed genome rearrangements in the *de novo* acquisition of AR [22]. However, their study did not reveal the mechanism how the rearrangements could lead to AR. Since most antibiotics induces ROS, this mechanism is sufficient for the survival in most cases.

To be noted, multiple AR mechanisms are coordinated and are involved sequentially in a stepwise manner. At the early stage, the tRNA up-regulation provided a prompt protection at early stage. From the 8^th^ generation, the efflux pump expression was increased. From the 22^nd^ generation, the genomic mutation provided effective resistance at late stage (Fig. 6B). This coincides with the time-scale of the regulation happens at different levels: the translation regulates most quickly; the transcription reacts slower, and the genomic mutations usually need prolonged time to accumulate. However, the effect size is *vice versa*. We previously reported that the tRNA up-regulation efficiently counteracts the ROS generated by hydrogen peroxide within 90 minutes by maintaining translation process [12], demonstrating that the translational response can provide the initial protection against antibiotics.

Although the excessive tRNA provides advantages in the presence of antibiotics, they have to be tightly controlled in the absence of antibiotics. Under normal conditions, excessive tRNA reduces the translational pausing and leads to massive misfolding, creating additional stress [13, 14] and thus reduces the fitness [12]. In our competition experiment, the tRNA elevated strain showed disadvantage under normal conditions. However, in the presence of antibiotics, the ROS slows down the translation in general. Therefore, elevating tRNA helps to maintain translational elongation rate is beneficial for the survival [12, 21].

Interestingly, the tRNA genes and their flanking sequences seems to be sensitive to the DNA damage and recombination. It has been shown that the tRNA genes serve as high frequency integration sites for genetic elements in prokaryote genomes [20]. The highly similar and conserved sequences and small size favors the recognition of integration sites. Also, these regions are also favorable in horizontal gene transfer [23]. However, these previous studies did not investigate the recombination around the tRNA genes and the influence of tRNA expression. Maintaining the translational elongation rate does not require overexpression of all tRNAs. This largely increases the probability of successful “resistance recombination”. Random elevation of several tRNAs is enough to provide survival advantage in the presence of antibiotics, which has been validated using the pRIL plasmid (Fig. 3D). Moreover, it has been reported that the bacterial tRNA genes are neither evenly nor randomly distributed in genomes [24, 25]. This distribution pattern might be also linked to the high frequency of recombination near tRNA genes.

Finally, our study provided a hint to suppress the evolution of intrinsic AR: suppressing the bacterial recombination systems while applying antibiotics. This can effectively delay or even abolish the emergence of resistance against most antibiotics. Since RecA is the most important recombinase, and it is highly conserved in almost all eubacteria, both in sequence and in structure [26], a small molecule which binds and blocks RecA of many bacterial species might be feasible. Although the risk of acquired AR by horizontal transfer or resistance genes still exists, this reduces the risk of intrinsic resistance, especially when using a newly artificial antibiotics, for which the resistance gene does not yet exist.

